# A computational framework to identify metabolic engineering strategies for the co-production of metabolites

**DOI:** 10.1101/2021.09.18.460904

**Authors:** Lavanya Raajaraam, Karthik Raman

## Abstract

Microbial production of chemicals is a more sustainable alternative to traditional chemical processes. However, the shift to bioprocess is usually accompanied by a drop in economic feasibility. Co-production of more than one chemical can improve the economy of bioprocesses, enhance carbon utilization and also ensure better exploitation of resources. While a number of tools exist for *in silico* metabolic engineering, there is a dearth of computational tools that can co-optimize the production of multiple metabolites. In this work, we propose an eXtended version of Flux Scanning based on Enforced Objective Flux (XFSEOF), identify intervention strategies to co-optimize for a set of metabolites. XFSEOF can be used to identify all pairs of products that can be co-optimized with ease, by a single intervention. Beyond this, it can also identify higher-order intervention strategies for a given set of metabolites. We have employed this tool on the genome-scale metabolic models of *Escherichia coli* and *Saccharomyces cerevisiae*, and identified intervention targets that can co-optimize the production of pairs of metabolites under both aerobic and anaerobic conditions. Anaerobic conditions were found to support the co-production of a higher number of metabolites when compared to aerobic conditions in both organisms. The proposed computational framework will enhance the ease of study of metabolite co-production and thereby aid the design of better bioprocesses.

## 1 Introduction

Recent years have seen several advances in the usage of bioprocessing to produce a wide range of chemicals (1). Microorganisms can produce diverse and complex products from simple carbon sources. Nevertheless, there are many challenges in designing economically feasible bioprocesses. The advancements in synthetic biology have enabled the metabolic engineering of organisms to improve yield and productivity (2). Various computational strain design algorithms have been developed to identify the genetic manipulations required to over-produce a single product (3–5). Despite the increase in yield achieved through such rational strain design, the bioprocesses are unable to compete with the traditional chemical processes in many cases (6). This is due to two main reasons – (i) the cost of raw materials (ii) the maximum yield achievable for a given product in a given organism and environment is limited by the number of genetic manipulations that can be successfully implemented in a single strain (7). The cost of raw materials can be reduced by using agricultural waste as feedstock instead of a synthetic nutrient medium. The latter can be overcome by co-producing multiple products in the same bioprocess (8).

Co-production equips us to exploit the system in a better fashion and produce more valuable products from the same raw materials. A high-value, low-volume chemical can be co-produced with a low-value, high-volume product in order to increase the economic feasibility, as in the case of riboflavin and butanol, respectively (6). Co-production is also beneficial when we need to co-optimize a cocktail of metabolites rather than a single metabolite, as in the case of biofuels and fatty acids (9). It can also balance the carbon metabolism, as in the case of uridine and acetoin (10). High carbon inflow towards uridine causes excess production of acetate, which hampers the growth of the organism. Conversion of acetate to acetoin prevents over-acidification of the nutrient medium and thereby improves growth and uridine production. There are many studies that have successfully achieved co-production of a variety of products with/without genetic manipulation of the organisms. Polyhydroxyalkanoates are one of the common class of metabolites that are co-produced with other metabolites (11–13). Butanol and hydrogen have been co-produced in *Clostridium beijerinckii* (14), and ethanol and xylitol have been co-produced in *Candida tropicalis* (15,16). The carbon source, nutrient medium, pH etc., are optimized in such cases to improve the yield of metabolites. Metabolic engineering can further expand the number of products that are co-produced and also improve their yield significantly. Multiple metabolites like ethanol, isopropanol, butanol and 2,3 butanediol have been co-produced by optimizing the acetone-butanol-ethanol (ABE) fermentation pathway in *Clostridium acetobutylicum* (17). Nisin and 3-phenyllactic acid, two antimicrobial agents, were co-produced in *Lactococcus lactis* through genetic manipulation (18). Non-native metabolites can also be co-produced with other metabolites, as in the case of butanol and riboflavin, by engineering the heterologous pathway in *C. acetobutylicum* (6).

Although many strain design algorithms have been successfully employed for metabolically engineering organisms to optimize a single product (19,20), few studies have applied it for co-production. The studies listed above only use existing literature and readily apparent deletion targets to achieve co-production. This limits the robustness of the bioprocesses that are designed. There is a lack of algorithms that can be easily applied to study co-production. In this study, we have extended the Flux Scanning based on Enforced Objective Flux (FSEOF) (21) algorithm to study co-production. Further, while deletion targets can be obtained for metabolites independently using existing algorithms like OptKnock (3), OptGene (4), there are very few algorithms that can identify amplification targets (22). In order to identify amplification targets in addition to knock-out targets for co-optimizing a set of metabolites, we propose a new methodology, eXtended (XFSEOF), adapting the FSEOF algorithm. XFSEOF has a simple computational framework that can be easily modified, and it also provides the entire set of potential intervention strategies in a single run while many algorithms are sequential, returning one intervention target per run. The utility of the potential intervention strategies obtained was further assessed using Flux Variability Analysis (FVA). We applied XFSEOF to evaluate all possible pairs of secretory metabolites in *Escherichia coli* and *Saccharomyces cerevisiae*. The different pairs of metabolites that can be co-produced through a single reaction deletion or amplification were obtained. This analysis helps us choose favorable pairs of metabolites for which higher-order intervention strategies can be obtained. We have demonstrated this by identifying the amplification targets, knock-out targets and mixed intervention strategies of size up to three to co-optimize the production of isobutanol and succinic acid in *S. cerevisiae*. Higher-order intervention strategies were able to achieve better yield with very little reduction in growth rate. Overall, our analyses provide an overall picture of the biosynthetic capabilities of an organism, particularly highlighting key interdependencies in metabolism.

## 2 Methods

### 2.1 Flux Balance Analysis (FBA)

FBA is a widely used steady-state constraint-based modelling approach to predict the metabolic capabilities of a variety of organisms (23–25). The metabolic network of an organism, which comprises all reactions known to occur in the organism, is represented as a stoichiometric matrix *S*, of size *m* × *n*, where *m* is the number of metabolites and *n* is the number of reactions (Fig 1a). The entries in the *j*^th^ column of *S* represent the stoichiometric coefficients of the metabolites that participate in the *j*^th^ reaction. The minimum and maximum values of flux that any reaction can assume are constrained by the lower and upper bounds, respectively. The flux through a reaction under a given set of conditions, at steady-state, is calculated by solving a linear programming (LP) problem. The LP problem is formulated as:

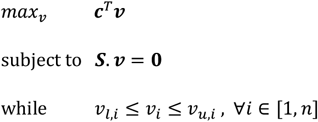

where *c* is a vector of weights denoting the contribution of each of the *n* reactions to the objective function, *v* ∊ *R^n^* is the vector of metabolic fluxes, ***v_l_*** and ***v_u_*** are vectors representing the lower and upper bounds for the reaction fluxes, respectively (25).

**Figure 1.**
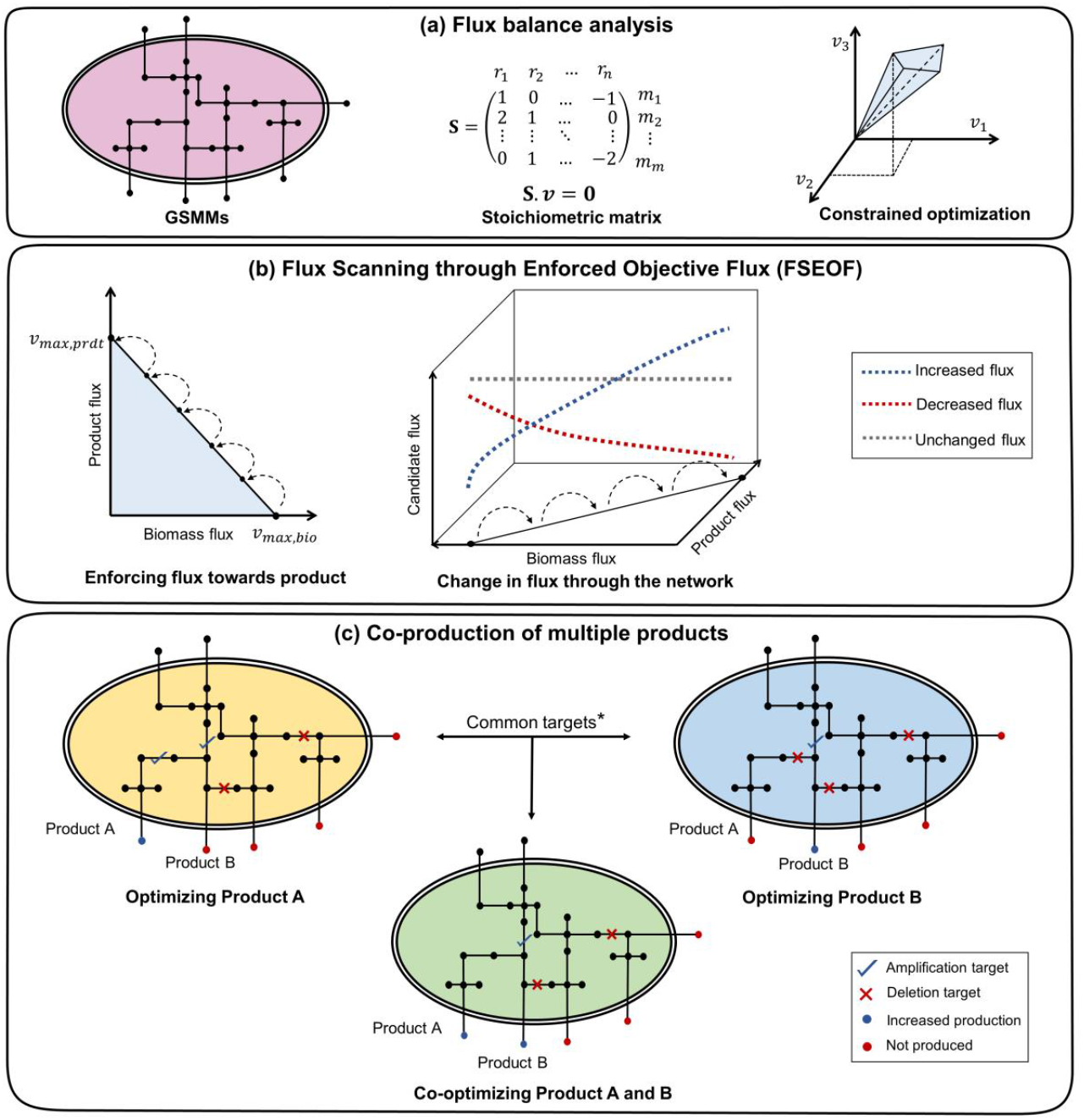
Workflow for XFSEOF. (a) The GSMM is represented as a stoichiometric matrix, which is used for FBA. (b) The flux through the product is increased in steps, and flux changes through all other reactions are studied. The reactions that have increased flux with an increase in product flux are potential amplification targets. The reactions that have decreased flux are potential deletion targets, while those with unchanged or oscillatory fluxes are excluded. (c) The targets common to products A and B are the potential targets for co-optimization. *The union of all potential targets for products A and B is used for higher-order intervention strategies

### 2.2 Flux Variability Analysis (FVA)

FVA is used to identify the range of fluxes of each reaction that still satisfy the constraints, where two optimization problems are solved for each flux *v_i_* of interest.

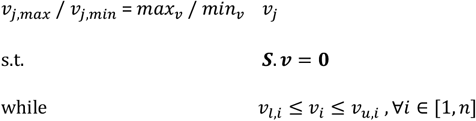

where *v* ∊ *R^n^* is the vector of metabolic fluxes, *v_j,max_* and *v_j,min_* are the maximum and minimum values of fluxes, respectively for each reaction flux *v_j_* (26).

### 2.3 Flux Scanning based on Enforced Objective Flux (FSEOF)

FSEOF (21) is a method used to identify potential reaction deletion and amplification targets in metabolic networks by observing the change in the reaction fluxes when the system moves from the wild-type flux of the target product to the theoretical maximum flux of the product (Fig 1b). The maximum biomass *v_max,bio_* and maximum product *v_max,prdt_* fluxes are obtained by performing FBA with the biomass reaction and the exchange reaction of the product as the objective, respectively. The flux of the target reaction, *v_prdt_* is pinned to *x*% *of v_max,prdt_* (*x* = 0 → 100). The change in the flux of a reaction, *v_j_*, is studied as the product flux, *v_prdt_* is increased, and it is classified as a potential deletion or amplification target based on the decrease or increase in its flux, respectively. The reactions that undergo no change or oscillations in the fluxes are discarded from the set of potential intervention strategies. The set of potential intervention strategies obtained are assessed by simulating each intervention and performing FVA on the mutant.

### 2.4 XFSEOF: Extending FSEOF for co-optimization

The Genome-Scale Metabolic Models (GSMMs) of *E. coli* iML1515 and *S. cerevisiae* iMM904 were obtained from the BiGG models database (http://bigg.ucsd.edu/) (15). The simulations were done with the following constraints on uptakes: −10 mmol/gDW/h glucose and −2 mmol/gDW/h oxygen for aerobic conditions and −10 mmol/gDW/h glucose and zero oxygen uptake for anaerobic conditions. The potential intervention strategies for all secretory metabolites (metabolites that can be secreted into the medium) in the organism were obtained as described in Section 2.3. Exchange and transport reactions were removed from the list of potential intervention strategies to increase the relevance of the results and reduce the computational time. The standard intervention strategies were obtained for all possible pairs of secretory metabolites.

The reliability of the potential deletion targets was studied by deleting the reaction and performing FVA for the product reaction. The reliability of the potential amplification targets was studied by fixing the flux bound of the amplification target to its theoretical maximum and performing FVA for the product reaction. Any reaction with more than a 5% increase in the maximum product flux and less than 75% decrease in biomass flux is considered a promising intervention strategy. FVA was performed using fastFVA to reduce the computational time (26). To obtain higher-order intervention strategies, the potential targets obtained earlier for a given set of metabolites were combined, and all possible combinations of intervention strategies of a certain size (up to three) were evaluated using FVA. The score for each product *i* (total number of products, *n*) and the overall score are calculated as

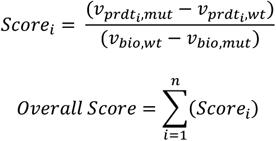

where *v_prdt_i_,mut_* and *v_prdt_i_,wt_* are the mutant and wild-type maximum fluxes of the exchange reaction of the product *i*, and *v_bio,mut_* and *v_bio,wt_* are the mutant and wild-type fluxes of the biomass reaction. *Score_i_* denotes the score for the individual product while *overall score* denotes the cumulative score for the set of metabolites. All simulations were performed in MATLAB R2018a (MathWorks Inc., USA) using the COBRA Toolbox v3.0 (27) and IBM ILOG CPLEX 12.8 as the linear programming solver.

## 3 Results

Metabolic engineering strategies for the co-production of all pairs of secreted metabolites in *E. coli* and *S. cerevisiae* were obtained using XFSEOF as described in Section 2.4. We identified the intervention strategies required to optimize the co-production of metabolites in both aerobic and anaerobic conditions. Anaerobic conditions favor the co-production of more pairs of metabolites when compared to aerobic conditions. The intervention strategy for each pair of metabolites is scored as in Section 2.4. The best intervention strategy can be chosen using the *overall score*. In cases where one metabolite might be favored over the others, the individual scores, *Score_i_* can be used to choose the best intervention strategies. Some of the intervention strategies obtained have been successfully verified through experimental studies in literature. This shows the credibility of the intervention strategies obtained. We discuss a few of the industrially significant metabolites and their intervention strategies, along with supporting literature. We also propose many other intervention strategies, which form a ready short-list for experimental validation. We were able to identify other hitherto unexplored intervention strategies, which may be better alternatives to those in existing literature, further demonstrating the utility of the algorithm.

### 3.1 Co-production in *Escherichia coli*

*E. coli* is one of the well-studied model organisms and has high-quality GSMMs available. The latest GSMM, *i*ML1515 (28), was used in this study, and the co-production of 337 secretory metabolites was studied in both aerobic and anaerobic conditions.

#### 3.1.1 Aerobic fermentation

Co-production of all pairs of metabolites was studied in *E. coli* using XFSEOF and FVA as described in Sections 2.3 and 2.4. Out of ^337^*C*_2_ pairs of secretory metabolites, only 237 could be successfully overproduced through deletion or amplification of a single reaction. The intervention strategies for a few industrially significant pairs of metabolites are listed in Table 1. One of the important pairs of metabolites that can be easily co-produced is L-lysine, a food additive and drug additive and cadaverine, which is essential for polyamide production. XFSEOF was able to identify several reactions from the diaminopimelate pathway (DAP), which can be over-expressed to co-produce L-lysine and cadaverine. An experimental study by Xu *et al*. (29) demonstrates the effect of engineering the DAP pathway in *E. coli* for the production of L-lysine. This indicates the reliability of the results obtained through computational methods. Another significant result is the co-production of succinate and ethanol through the amplification of glyceraldehyde-3-phosphate dehydrogenase. Other studies have successfully co-produced ethanol and succinate by other genetic manipulations (30).

**Table 1.**
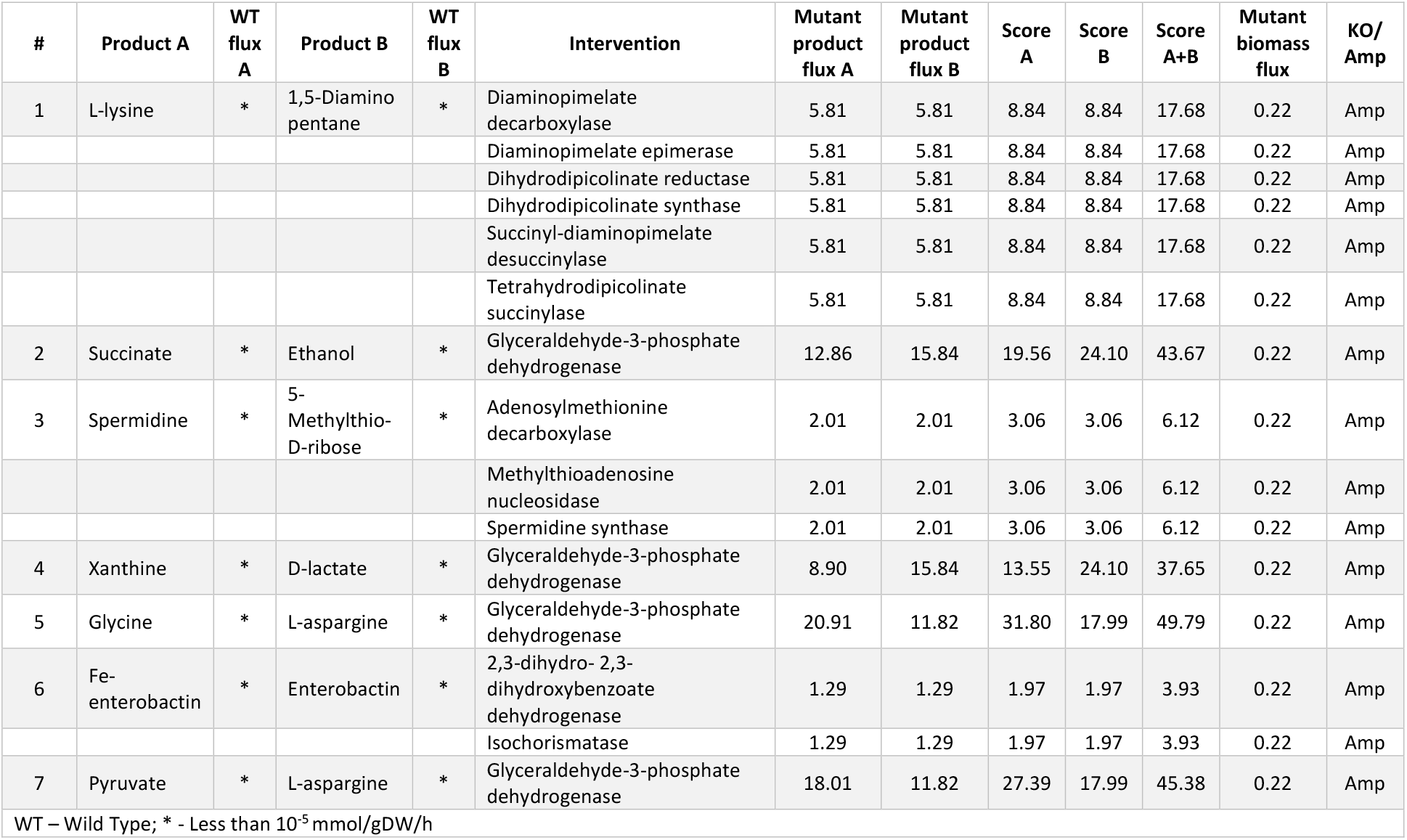
Intervention strategies for co-production of pairs of metabolites in *E. coli* under aerobic conditions

#### 3.1.2 Anaerobic fermentation

Anaerobic conditions support the co-production of more metabolites when compared to aerobic conditions. More than 1000 pairs of metabolites can be co-produced, out of which few are listed in Table 2. L-lysine and cadaverine can be co-produced under anaerobic conditions too. But the maximum flux achievable is lower when compared to aerobic conditions. The yield of metabolites like acetate, formate and hexanoate can be co-optimized by deleting acetaldehyde dehydrogenase or alcohol dehydrogenase. We also found that succinate and lactate can be co-produced by the knock-out of pyruvate formate lyase. The effect of deletion of *pflB* gene encoding pyruvate formate lyase has been experimentally verified in *E. coli* for succinate production (31) and lactate production (32) through separate studies. This shows that there are multiple co-production strategies available in the existing literature that can be easily utilized to design an efficient process.

**Table 2.** Intervention strategies for co-production of pairs of metabolites in *E. coli* under anaerobic conditions

| # | Product A | WT flux A | Product B | WT flux B | Intervention | Mutant product flux A | Mutant product flux B | Mutant biomass flux | Score A | Score B | Score A+B | KO/Amp |
| --- | --- | --- | --- | --- | --- | --- | --- | --- | --- | --- | --- | --- |
| 1 | Acetate | 8.83 | Formate | 18.22 | Acetaldehyde dehydrogenase | 18.18 | 36.79 | 0.12 | 237.95 | 472.31 | 710.26 | KO |
|  |  |  |  |  | Alcohol dehydrogenase | 18.18 | 36.79 | 0.12 | 237.95 | 472.31 | 710.26 | KO |
| 2 | Succinate | 0.05 | D-lactate | 3.76E-04 | Glyceraldehyde-3-phosphate dehydrogenase | 14.98 | 19.41 | 0.04 | 126.78 | 164.92 | 291.71 | Amp |
|  |  |  |  |  | Acetaldehyde dehydrogenase | 9.07 | 18.12 | 0.12 | 229.50 | 461.04 | 690.55 | KO |
|  |  |  |  |  | Alcohol dehydrogenase | 9.07 | 18.12 | 0.12 | 229.50 | 461.04 | 690.55 | KO |
|  |  |  |  |  | Pyruvate formate lyase | 0.49 | 17.76 | 0.12 | 10.57 | 426.65 | 437.22 | KO |
| 3 | L-alanine | * | Xanthine | * | Glyceraldehyde-3-phosphate dehydrogenase | 13.56 | 0.97 | 0.04 | 115.18 | 8.23 | 123.41 | Amp |
| 4 | Spermidine | * | 5-Methylthio-D-ribose | * | Adenosylmethionine decarboxylase | 0.64 | 0.64 | 0.04 | 5.46 | 5.46 | 10.93 | Amp |
|  |  |  |  |  | Methylthioadenosine nucleosidase | 0.64 | 0.64 | 0.04 | 5.46 | 5.46 | 10.93 | Amp |
|  |  |  |  |  | Spermidine synthase | 0.64 | 0.64 | 0.04 | 5.46 | 5.46 | 10.93 | Amp |
| 5 | L-aspartate | * | L-glutamate | * | Glyceraldehyde-3-phosphate dehydrogenase | 6.03 | 4.52 | 0.04 | 51.19 | 38.39 | 89.59 |  |
| 6 | L-lysine | * | 1,5-Diamino pentane | * | Diaminopimelate decarboxylase | 2.72 | 2.72 | 0.04 | 23.04 | 23.04 | 46.07 | Amp |
|  |  |  |  |  | Diaminopimelate epimerase | 2.72 | 2.72 | 0.04 | 23.04 | 23.04 | 46.07 | Amp |
|  |  |  |  |  | Dihydrodipicolinate synthase | 2.72 | 2.72 | 0.04 | 23.04 | 23.04 | 46.07 | Amp |
|  |  |  |  |  | Succinyl-diaminopimelate desuccinylase | 2.72 | 2.72 | 0.04 | 23.04 | 23.04 | 46.07 | Amp |
|  |  |  |  |  | Tetrahydrodipicolinate succinylase | 2.72 | 2.72 | 0.04 | 23.04 | 23.04 | 46.07 | Amp |
| 7 | Acetate | 8.83 | Hexanoate | * | Acetaldehyde dehydrogenase | 18.18 | 4.53 | 0.12 | 237.95 | 115.26 | 353.21 | KO |
|  |  |  |  |  | Alcohol dehydrogenase | 18.18 | 4.53 | 0.12 | 237.95 | 115.26 | 353.21 | KO |
| WT – Wild Type; * - Less than 10<sup>-5</sup> mmol/gDW/h |  |  |  |  |  |  |  |  |  |  |  |  |

### 3.2 Co-production in *Saccharomyces cerevisiae*

Another industrially relevant and well-studied model organism is *S. cerevisiae*. Though heterologous pathways have not been analyzed in this study, one can easily modify the GSMM and apply XFSEOF to identify co-production strategies for heterologous metabolites. Since *S. cerevisiae* is a better candidate for recombinant protein production, it is essential to study co-production in yeast (33). It can also produce more complex metabolites when compared to *E. coli* and is, therefore, a favorable candidate for bio-production. The latest GSMM iMM904 (34) was used, and the ability to optimize the co-production of 164 secretory metabolites was studied in both aerobic and anaerobic conditions.

#### 3.2.1 Aerobic fermentation

We found that many industrially important metabolites like ethanol and L-alanine, and 4-aminobutanoate and L-serine can be co-produced in *S. cerevisiae* under aerobic conditions. We were also able to co-optimize isobutyl alcohol and 2-methyl propanal, which are long-chain alcohols that are used as biofuels. The deletion of pyruvate dehydrogenase increases the production of pyruvate and acetate, as shown in Table 3. Although the deletion of pyruvate dehydrogenase has not been experimentally verified as yet, a similar study has been carried out in *E. coli* (35). In this study, it has been shown that the deletion of the genes encoding pyruvate dehydrogenase improves pyruvate production (35). In addition to pyruvate dehydrogenase, XFSEOF was able to identify several other amplification targets, which can also improve the production of pyruvate and acetate.

**Table 3.** Intervention strategies for co-production of pairs of metabolites in *S. cerevisiae* under aerobic conditions

| # | Product A | WT flux A | Product B | WT flux B | Intervention | Mutant product flux A | Mutant product flux B | Mutant biomass flux | Score A | Score B | Score A+B | KO/ Amp |
| --- | --- | --- | --- | --- | --- | --- | --- | --- | --- | --- | --- | --- |
| 1 | Ethanol | 15.81 | L-alanine | 1.69E-04 | Sedoheptulose 1,7-bisphosphate D-glyceraldehyde-3-phosphate-lyase | 18.45 | 0.15 | 0.07 | 12.22 | 0.71 | 12.93 | Amp |
|  |  |  |  |  | Phosphofructokinase (s7p) | 18.45 | 0.15 | 0.07 | 12.22 | 0.71 | 12.93 | Amp |
| 2 | Acetate | 3.55E-03 | 2,3 butanediol | 3.2E-04 | Pyruvate dehydrogenase | 1.48 | 0.25 | 0.28 | 230.94 | 38.48 | 269.42 | KO |
|  |  |  |  |  | Enolase | 10.71 | 10.27 | 0.07 | 49.61 | 47.61 | 97.22 | Amp |
|  |  |  |  |  | Fructose-bisphosphate aldolase | 10.70 | 10.27 | 0.07 | 49.62 | 47.64 | 97.26 | Amp |
|  |  |  |  |  | Glyceraldehyde-3-phosphate dehydrogenase | 10.71 | 10.27 | 0.07 | 49.61 | 47.61 | 97.22 | Amp |
|  |  |  |  |  | Triose-phosphate isomerase | 10.70 | 10.27 | 0.07 | 49.62 | 47.64 | 97.26 | Amp |
| 3 | Isobutyl alcohol | * | Succinate | * | Pyruvate decarboxylase | 7.63 | 5.45 | 0.19 | 79.29 | 56.66 | 135.95 | KO |
|  |  |  |  |  | Enolase | 9.56 | 12.97 | 0.07 | 44.30 | 60.10 | 104.40 | Amp |
|  |  |  |  |  | Glyceraldehyde-3-phosphate dehydrogenase | 9.56 | 12.97 | 0.07 | 44.30 | 60.13 | 104.43 | Amp |
|  |  |  |  |  | Triose-phosphate isomerase | 9.56 | 12.97 | 0.07 | 44.33 | 60.17 | 104.50 | Amp |
| 4 | 2-methylpropanal | * | Isobutyl alcohol | * | 3 Methyl 2 oxobutanoate decarboxylase | 6.50 | 9.36 | 0.07 | 30.14 | 43.39 | 73.54 | Amp |
|  |  |  |  |  | Acetolactate synthase mitochondrial | 6.50 | 9.36 | 0.07 | 30.14 | 43.40 | 73.54 | Amp |
|  |  |  |  |  | Dihydroxy acid dehydratase 2 3 dihydroxy 3 methylbutanoate mitochondrial | 6.50 | 9.36 | 0.07 | 30.14 | 43.40 | 73.54 | Amp |
|  |  |  |  |  | Enolase | 8.95 | 9.56 | 0.07 | 41.47 | 44.30 | 85.77 | Amp |
|  |  |  |  |  | Glyceraldehyde-3-phosphate dehydrogenase | 8.95 | 9.56 | 0.07 | 41.47 | 44.30 | 85.77 | Amp |
|  |  |  |  |  | Acetohydroxy acid isomeroreductase mitochondrial | 6.50 | 9.36 | 0.07 | 30.14 | 43.40 | 73.54 | Amp |
|  |  |  |  |  | Triose-phosphate isomerase | 8.94 | 9.56 | 0.07 | 41.49 | 44.33 | 85.82 | Amp |
| 5 | L-Glutamate | * | 2-Oxoglutarate | * | Citrate synthase | 2.93 | 1.95 | 0.07 | 13.58 | 9.05 | 22.63 | Amp |
|  |  |  |  |  | Enolase | 4.50 | 4.04 | 0.07 | 20.85 | 18.73 | 39.58 | Amp |
|  |  |  |  |  | Fructose-bisphosphate aldolase | 4.50 | 4.04 | 0.07 | 20.86 | 18.73 | 39.60 | Amp |
|  |  |  |  |  | Glyceraldehyde-3-phosphate dehydrogenase | 4.50 | 4.04 | 0.07 | 20.85 | 18.73 | 39.58 | Amp |
|  |  |  |  |  | Isocitrate dehydrogenase | 2.99 | 2.00 | 0.07 | 13.87 | 9.25 | 23.12 | Amp |
|  |  |  |  |  | Triose-phosphate isomerase | 4.50 | 4.04 | 0.07 | 20.86 | 18.73 | 39.60 | Amp |
| 6 | Acetate | 3.55E-03 | Pyruvate | 3.24E-04 | Pyruvate dehydrogenase | 1.48 | 0.25 | 0.28 | 230.94 | 38.48 | 269.42 | KO |
|  |  |  |  |  | Aspartate-semialdehyde dehydrogenase | 8.79 | 11.85 | 0.07 | 41.14 | 55.49 | 96.63 | Amp |
|  |  |  |  |  | Enolase | 10.71 | 14.42 | 0.07 | 49.61 | 66.83 | 116.44 | Amp |
|  |  |  |  |  | Fructose-bisphosphate aldolase | 10.70 | 14.41 | 0.07 | 49.62 | 66.85 | 116.47 | Amp |
|  |  |  |  |  | Glyceraldehyde-3-phosphate dehydrogenase | 10.71 | 14.42 | 0.07 | 49.61 | 66.83 | 116.44 | Amp |
|  |  |  |  |  | Triose-phosphate isomerase | 10.70 | 14.41 | 0.07 | 49.62 | 66.85 | 116.47 | Amp |
| 7 | 4-Aminobutanoate | * | L-serine | * | Glyceraldehyde-3-phosphate dehydrogenase | 4.77 | 6.98 | 0.07 | 22.10 | 32.34 | 54.44 | Amp |
|  |  |  |  |  | Triose-phosphate isomerase | 4.77 | 6.97 | 0.07 | 22.11 | 32.35 | 54.45 | Amp |
| 8 | L-alanine | 1.69E-04 | L-cysteine | * | Glucose 6-phosphate dehydrogenase | 10.60 | 1.20 | 0.07 | 49.11 | 5.56 | 54.67 | Amp |
|  |  |  |  |  | Phosphogluconate dehydrogenase | 10.60 | 1.20 | 0.07 | 49.11 | 5.56 | 54.67 | Amp |
|  |  |  |  |  | 6-phosphogluconolactonase | 10.60 | 1.20 | 0.07 | 49.11 | 5.56 | 54.67 | Amp |
|  |  |  |  |  | Ribulose 5-phosphate 3-epimerase | 10.60 | 1.20 | 0.07 | 49.11 | 5.56 | 54.67 | Amp |
|  |  |  |  |  | Transketolase | 10.60 | 1.20 | 0.07 | 49.12 | 5.56 | 54.68 | Amp |
|  |  |  |  |  | Ribose-5-phosphate isomerase | 18.16 | 2.21 | 0.08 | 85.44 | 10.40 | 95.84 | Amp |
| 9 | Sn-Glycero-3-phosphocholine | * | L-methionine | * | Methionine synthase | 0.02 | 2.19 | 0.07 | 0.11 | 10.17 | 10.27 | Amp |
|  |  |  |  |  | 5 10 methylene-tetrahydrofolate reductase NADPH | 0.02 | 2.19 | 0.07 | 0.11 | 10.17 | 10.27 | Amp |
|  |  |  |  |  | Ribose-5-phosphate isomerase | 0.66 | 1.71 | 0.08 | 3.11 | 8.03 | 11.14 | Amp |
| 10 | 2 methylbutyl acetate | * | 2-Methyl-1-butanol | * | Ribose-5-phosphate isomerase | 3.00 | 4.42 | 0.08 | 14.13 | 20.78 | 34.92 | Amp |
| WT – Wild Type; * - Less than 10<sup>-5</sup> mmol/gDW/h |  |  |  |  |  |  |  |  |  |  |  |  |

#### 3.2.2 Anaerobic fermentation

As in the case of *E. coli*, anaerobic fermentation enables the co-production of more pairs of metabolites in *S. cerevisiae* when compared to aerobic fermentation. 2-methyl-1-butanol, which is an important solvent used in the manufacture of pesticides and paints and isobutyl alcohol, which is a biofuel, can be co-produced by the amplification of malic enzyme, as shown in Table 4. Formate, which is used in dyeing and printing, can be co-produced with spermidine, a metabolite increasingly studied for its anti-ageing properties (36), through the amplification of a number of reactions. These strategies not only include readily apparent reactions that are involved in spermidine synthesis like spermidine synthase and adenosylmethionine decarboxylase but also provide some non-intuitive strategies like the amplification of aspartate transaminase or 2-keto-4-methylthiobutyrate transaminase. We also found that the deletion of pyruvate decarboxylase improves the production of succinate, isobutyl alcohol and pyruvate. The effect of deletion of pyruvate decarboxylase has been studied in *S. cerevisiae*, and the improvement in the production of pyruvate (37) and succinate (38) has been verified experimentally in separate studies in literature.

**Table 4.** Intervention strategies for co-production of pairs of metabolites in *S. cerevisiae* under anaerobic conditions

| # | Product A | WT flux A | Product B | WT flux B | Intervention | Mutant product flux A | Mutant product flux B | Mutant biomass flux | Score A | Score B | Score A+B | KO/ Amp |
| --- | --- | --- | --- | --- | --- | --- | --- | --- | --- | --- | --- | --- |
| 1 | 2-Methyl-1-butanol | * | Isobutyl alcohol | * | Malic enzyme NADP mitochondrial | 0.03 | 9.64 | 0.05 | 0.20 | 61.30 | 61.51 | Amp |
| 2 | Isobutyl alcohol | * | Pyruvate | * | Pyruvate decarboxylase | 8.73 | 5.66 | 0.11 | 88.74 | 57.57 | 146.31 | KO |
|  |  |  |  |  | Enolase | 9.68 | 9.97 | 0.05 | 61.59 | 63.45 | 125.03 | Amp |
|  |  |  |  |  | Fructose-bisphosphate aldolase | 9.68 | 9.96 | 0.05 | 61.67 | 63.47 | 125.14 | Amp |
|  |  |  |  |  | Glyceraldehyde-3-phosphate dehydrogenase | 9.68 | 9.97 | 0.05 | 61.60 | 63.45 | 125.05 | Amp |
|  |  |  |  |  | Triose-phosphate isomerase | 9.68 | 9.96 | 0.05 | 61.66 | 63.47 | 125.13 | Amp |
| 3 | Formate | 6.26E-04 | Spermidine | * | Adenosylmethionine decarboxylase | 1.22 | 1.22 | 0.05 | 7.76 | 7.76 | 15.52 | Amp |
|  |  |  |  |  | Aspartate transaminase | 1.21 | 1.21 | 0.05 | 7.75 | 7.76 | 15.51 | Amp |
|  |  |  |  |  | 2 3 diketo 5 methylthio 1 phosphopentane degradation | 1.22 | 1.22 | 0.05 | 7.76 | 7.76 | 15.52 | Amp |
|  |  |  |  |  | 5-Methylthio-5-deoxy-D-ribulose 1-phosphate dehydratase | 1.22 | 1.22 | 0.05 | 7.76 | 7.76 | 15.52 | Amp |
|  |  |  |  |  | 5 methylthioadenosine phosphorylase | 1.22 | 1.22 | 0.05 | 7.76 | 7.76 | 15.52 | Amp |
|  |  |  |  |  | 5-methylthioribose-1-phosphate isomerase | 1.22 | 1.22 | 0.05 | 7.76 | 7.76 | 15.52 | Amp |
|  |  |  |  |  | Spermidine synthase | 1.22 | 1.22 | 0.05 | 7.76 | 7.76 | 15.52 | Amp |
|  |  |  |  |  | 2-keto-4-methylthiobutyrate transamination | 1.22 | 1.22 | 0.05 | 7.76 | 7.76 | 15.52 | Amp |
| 4 | 4-Amino butanoate | * | Isobutyl acetate | * | Enolase | 2.84 | 2.91 | 0.05 | 18.04 | 18.50 | 36.53 | Amp |
|  |  |  |  |  | Fructose-bisphosphate aldolase | 2.84 | 2.90 | 0.05 | 18.11 | 18.50 | 36.61 | Amp |
|  |  |  |  |  | Glyceraldehyde-3-phosphate dehydrogenase | 2.85 | 2.91 | 0.05 | 18.11 | 18.50 | 36.61 | Amp |
|  |  |  |  |  | Triose-phosphate isomerase | 2.84 | 2.90 | 0.05 | 18.11 | 18.50 | 36.61 | Amp |
| 5 | 2-Methyl-1-butanol | * | Glycine | * | Aspartate kinase | 0.05 | 4.55 | 0.05 | 0.32 | 28.95 | 29.26 | Amp |
|  |  |  |  |  | Glucose 6-phosphate dehydrogenase | 0.08 | 4.15 | 0.05 | 0.52 | 26.39 | 26.91 | Amp |
|  |  |  |  |  | Phosphogluconate dehydrogenase | 0.08 | 4.15 | 0.05 | 0.52 | 26.39 | 26.91 | Amp |
|  |  |  |  |  | Homoserine dehydrogenase NADH irreversible | 0.05 | 4.55 | 0.05 | 0.32 | 28.95 | 29.26 | Amp |
|  |  |  |  |  | Homoserine kinase | 0.05 | 4.55 | 0.05 | 0.32 | 28.91 | 29.22 | Amp |
|  |  |  |  |  | 6-phosphogluconolactonase | 0.08 | 4.15 | 0.05 | 0.52 | 26.39 | 26.91 | Amp |
|  |  |  |  |  | Ribulose 5-phosphate 3-epimerase | 0.08 | 4.15 | 0.05 | 0.52 | 26.39 | 26.91 | Amp |
|  |  |  |  |  | Ribose-5-phosphate isomerase | 3.31 | 4.01 | 0.06 | 21.52 | 26.04 | 47.56 | Amp |
|  |  |  |  |  | Threonine synthase | 0.05 | 4.55 | 0.05 | 0.32 | 28.91 | 29.22 | Amp |
|  |  |  |  |  | Transketolase | 0.08 | 4.15 | 0.05 | 0.52 | 26.39 | 26.91 | Amp |
|  |  |  |  |  | Transketolase | 0.08 | 4.15 | 0.05 | 0.52 | 26.39 | 26.91 | Amp |
| 6 | Isobutyl alcohol | * | Succinate | 0.67 | Pyruvate decarboxylase | 8.73 | 6.70 | 0.11 | 88.74 | 61.27 | 150.02 | KO |
| 7 | L-Glutamate | * | Xanthine | * | Fructose-bisphosphate aldolase | 2.81 | 0.95 | 0.05 | 17.90 | 6.04 | 23.94 | Amp |
|  |  |  |  |  | Glyceraldehyde-3-phosphate dehydrogenase | 2.81 | 0.95 | 0.05 | 17.90 | 6.04 | 23.93 | Amp |
|  |  |  |  |  | Triose-phosphate isomerase | 2.81 | 0.95 | 0.05 | 17.90 | 6.04 | 23.94 | Amp |
| 8 | Sorbitol | * | L-methionine | * | Ribose-5-phosphate isomerase | 5.61 | 1.28 | 0.06 | 36.45 | 8.29 | 44.74 | Amp |
| 9 | 2,3 butanediol | * | L-serine | * | Fructose-bisphosphate aldolase | 9.30 | 4.74 | 0.05 | 59.21 | 30.19 | 89.40 | Amp |
|  |  |  |  |  | Glyceraldehyde-3-phosphate dehydrogenase | 9.30 | 4.74 | 0.05 | 59.19 | 30.19 | 89.37 | Amp |
|  |  |  |  |  | Triose-phosphate isomerase | 9.30 | 4.74 | 0.05 | 59.21 | 30.19 | 89.40 | Amp |
| WT – Wild Type; * - Less than 10<sup>-5</sup> mmol/gDW/h |  |  |  |  |  |  |  |  |  |  |  |  |

#### 3.2.3 Higher-order intervention strategies – co-production of isobutanol and succinate

Higher-order intervention strategies can increase the maximum yield achievable for any product with a little difference in growth rate when compared to single interventions. But they are more cumbersome to identify, as the problem becomes time-consuming and computationally expensive. Instead of identifying higher-order targets for all metabolites in an organism, we have used the previous analysis to explore the metabolic capabilities of the organism and chose one set of metabolites to demonstrate the power of higher-order intervention strategies.

Isobutanol is a long-chain alcohol that is an attractive biofuel (39). Succinic acid is an important metabolite essential for the production of various other products like biodegradable polymers, fatty acids, butyrolactone and tetrahydrofuran (40). The co-production of isobutanol and succinate has been proposed as a sustainable and economical process by Xu *et al*., (41). They have discussed the development of various strains for the production of isobutanol and succinate separately. They emphasize how the co-production of isobutanol and succinate is not only of economic significance, but the high amount of carbon dioxide released from long-chain alcohol fermentation can be used for succinate production, and is hence also of ecological importance. But the article does not discuss any strategy to co-optimize the production of isobutanol and succinate.

Here we identified the higher-order intervention strategies (size up to three) for co-production of isobutanol and succinate in *S. cerevisiae* in aerobic conditions. More than 3700 interventions can improve the yield of both the metabolites when compared to the wild-type strain. Table 5 lists a few examples of each type of intervention strategy obtained, some of which are also found in experimental studies in literature. The deletion of pyruvate decarboxylase has been shown to improve the production of isobutanol by Kondo *et al*., (42). Zahoor *et al*., (38) have shown that both pyruvate decarboxylase deletion and fumarase deletion can increase the production of succinate. This shows the dependability of the results obtained using the algorithm.

**Table 5.** Higher-order intervention strategies for co-production of isobutanol and succinate in *S. cerevisiae* under aerobic conditions

| # | Intervention 1 | Intervention 2 | Intervention 3 | A/K/AK/AKK/AAK | Mutant flux 1 | Mutant flux 2 | Biomass flux | Score A | Score B | Score A+B |
| --- | --- | --- | --- | --- | --- | --- | --- | --- | --- | --- |
| 1 | Glyceraldehyde-3-phosphate dehydrogenase | Pyruvate kinase | NA | A | 7.19 | 8.60 | 0.20 | 81.73 | 97.87 | 179.59 |
| 2 | Enolase | Pyruvate kinase | NA | A | 7.13 | 8.60 | 0.20 | 81.08 | 97.87 | 178.95 |
| 3 | Glutamate 5-kinase | Phosphoglycerate dehydrogenase | Pyruvate decarboxylase | K | 7.96 | 5.80 | 0.19 | 79.81 | 58.12 | 137.93 |
| 4 | Fumarase | Phosphoserine phosphatase | Pyruvate decarboxylase | K | 7.95 | 5.78 | 0.19 | 79.78 | 58.05 | 137.84 |
| 5 | Aldehyde dehydrogenase | Glycerol 3 phosphate dehydrogenase | NA | AK | 2.99 | 2.52 | 0.22 | 43.76 | 36.77 | 80.54 |
| 6 | Succinate CoA ligase ADP forming | Pyruvate decarboxylase | NA | AK | 7.63 | 5.45 | 0.19 | 79.31 | 56.69 | 136.00 |
| 7 | Acetolactate synthase | Oxoglutarate dehydrogenase lipoamide | Guanylate kinase | AAK | 4.72 | 6.89 | 0.15 | 34.51 | 50.38 | 84.89 |
| 8 | Citrate synthase | Dihydroxy-acid dehydratase | Prephenate dehydrogenase | AAK | 4.73 | 6.98 | 0.14 | 32.31 | 47.71 | 80.02 |
| 9 | Aldehyde dehydrogenase | Ribonucleoside-diphosphate reductase | Ribulose 5-phosphate 3-epimerase | AKK | 2.99 | 2.48 | 0.22 | 43.84 | 36.32 | 80.16 |
| 10 | Succinate CoA ligase ADP forming | Phosphoserine phosphatase | Pyruvate decarboxylase | AKK | 7.95 | 5.78 | 0.19 | 79.78 | 58.05 | 137.84 |
| A- Amplification of all reactions; K- Knock-out of all reactions; AK – Amplification+ Knock-out; AAK – Amplification + Amplification + Knock-out; AKK – Amplification + Knock-out+ Knock-out |  |  |  |  |  |  |  |  |  |  |

## 4 Discussion

Chemical processes based on fossil fuels are cheaper when compared to bioprocesses, which leads to reluctance in the adoption of sustainable bioprocesses in industries. To improve the economic feasibility of a bioprocess, we can optimize the process variables and/or genetically engineer the microbes (43). Even then, in some cases, the bioprocess might be less lucrative when compared to their chemical counterparts (14). In such cases, we can co-produce multiple metabolites to improve the economic feasibility and efficiency of a bioprocess. Co-production also ensures better utilization of microbial capabilities, and better balance in the carbon metabolism. While there are multiple computational tools and algorithms to identify intervention strategies for a single product, there is a lack of readily appliable algorithms for co-production. As a result, almost no co-production study in existing literature was found to use computational algorithms to aid rational strain design. All of the studies rely on previous findings or readily apparent strategies to achieve co-production. This limits the intervention strategies designed.

In this study, we present XFSEOF, by adapting the effective FSEOF algorithm to study the co-optimization of a set of metabolites. FSEOF is a well-established constraint-based modelling algorithm, which has been used to reliably predict metabolic engineering strategies for a variety of systems (21,44–46). Extending FSEOF, we examined the co-production of multiple pairs of metabolites, and both deletion and amplification targets were obtained in *E. coli* and *S. cerevisiae* under both aerobic and anaerobic conditions. Anaerobic fermentation enabled the co-production of a higher number of metabolites when compared to aerobic fermentation in both organisms. This can be due to the incomplete respiration in the absence of oxygen that leads to the formation of multiple by-products. Also, *S. cerevisiae* produces more industrially significant metabolites when compared to *E. coli*. Some of these proposed intervention strategies have been verified experimentally by other studies in literature, as mentioned in Section 3. This shows the efficacy of the algorithm in furnishing reliable targets. XFSEOF also provides both intuitive and non-intuitive intervention strategies (as discussed in Section 3.2.2) for a set of metabolites, which exhibits the functionality of the algorithm.

The co-optimization analysis for all possible pairs of metabolites in the network is intended to be exploratory in order to give a larger picture of the metabolic capabilities of the organism. This analysis showed that around 200 pairs could be co-optimized in *E. coli* in aerobic conditions, and around 1000 pairs of metabolites could be co-optimized in all the other cases, respectively. Once a few sets of desirable metabolites are chosen from the previous analysis, XFSEOF can be extended to obtain higher-order intervention strategies, as shown in Section 3.2.3. Exploring the higher-order strategies can expand the efficiency of the intervention targets obtained. Since the evaluation of higher-order intervention strategies is laborious and computationally expensive, we have limited the size to a maximum of three manipulations at a time. It not only enhances yield, but also provides alternate routes to achieve a similar yield. The advantageous strategies can be chosen based on the ease of manipulation in an experimental setup in such cases.

The evaluation of the results is carried out using FVA, which ensures the robustness of the targets obtained. While FBA provides one optimal solution from the solution space, FVA gives the entire range of values the flux can take up. This is a significant difference that sets XFSEOF apart from other existing algorithms like OptKnock. Also, the algorithm validates and returns all the intervention strategies in a single run, contrary to the existing algorithms, most of which are sequential and require a separate run for each strategy obtained. The set of intervention strategies validated through FVA can be short-listed for experimental verification using the scores. The *overall score* can be used to compare the effectiveness of different intervention strategies. If one product is more favored than the others economically or otherwise, we can use the individual scores *Score_i_* to choose the appropriate strategy for the process that is formulated. The products can be chosen based on their economic value, or ease of co-production. One drawback of co-production is the cost associated with downstream processing. But this can be overcome by choosing easily separable products or choosing metabolites such that one is accumulated in the cell and one is secreted out, as in the case of polyhydroxy butyrate and succinate, respectively (47). XFSEOF not only identifies intervention strategies for co-production of a given set of metabolites, but allows us to explore the different combinations of products that can be co-produced in an easy and efficient manner.

## 5 Conclusion

Co-production can open new avenues for the sustainable production of chemicals. Designing bioprocesses for co-production using laboratory experiments alone is cumbersome and can result in sub-optimal strategies. XFSEOF empowers us to explore and exploit microbial systems in a better fashion. It can be used to computationally study and optimize co-production by identifying intervention strategies for multiple metabolites and thereby improve the efficiency of bioprocesses. It should be noted that the co-optimization analysis was limited to pairs of metabolites to reduce the computational time. But XFSEOF can be easily extended to co-optimize a more extensive set of metabolites. To conclude, this study can be used to identify various genetic manipulations that can co-optimize a set of products, which might be challenging to achieve through pure experimentation. It provides a novel and critical approach to study co-production computationally. We hope this study will aid the design and development of more sustainable bioprocesses.

## Data availability

All models used in this work and the codes used for our analysis are available at https://github.com/RamanLab/XFSEOF

## Acknowledgements

LR acknowledges the HTRA fellowship from the Ministry of Education, Government of India. KR acknowledges support from the Science and Engineering Board (SERB) MATRICS Grant MTR/2020/000490.

## Notes

http://bigg.ucsd.edu/

### Competing Interest Statement

The authors have declared no competing interest.

## References

1. Erickson B, Nelson, Winters P. Perspective on opportunities in industrial biotechnology in renewable chemicals. Biotechnology Journal. 2012;7(2):176–85.

2. Yadav B, Talan A, Tyagi RD, Drogui P. Concomitant production of value-added products with polyhydroxyalkanoate (PHA) synthesis: A review. Bioresource Technology. 2021 Oct;337:125419.

3. Burgard AP, Pharkya P, Maranas CD. Optknock: A bilevel programming framework for identifying gene strategies for microbial strain optimization. Biotechnol Bioeng. 2003 Dec 20;84(6):647–57.

4. Rocha I, Maia P, Rocha M, Ferreira EC. OptGene – a framework for in silico metabolic engineering. 2008;2.

5. Yang L, Cluett WR, Mahadevan R. EMILiO: A fast algorithm for genome-scale strain design. Metabolic Engineering. 2011 May;13(3):272–81.

6. Cai X, Bennett GN. Improving the Clostridium acetobutylicum butanol fermentation by engineering the strain for co-production of riboflavin. J Ind Microbiol Biotechnol. 2011 Aug 1;38(8):1013–25.

7. Silva F, Queiroz JA, Domingues FC. Evaluating metabolic stress and plasmid stability in plasmid DNA production by Escherichia coli. Biotechnology Advances. 2012 May 1;30(3):691–708.

8. da Silva TL, Gouveia L, Reis A. Integrated microbial processes for biofuels and high value-added products: the way to improve the cost effectiveness of biofuel production. Appl Microbiol Biotechnol. 2014 Feb;98(3):1043–53.

9. Xin F, Dong W, Jiang Y, Ma J, Zhang W, Wu H, et al. Recent advances on conversion and co-production of acetone-butanol-ethanol into high value-added bioproducts. Critical Reviews in Biotechnology. 2018 May 19;38(4):529–40.

10. Fan X, Wu H, Jia Z, Li G, Li Q, Chen N, et al. Metabolic engineering of Bacillus subtilis for the co-production of uridine and acetoin. Appl Microbiol Biotechnol. 2018 Oct;102(20):8753–62.

11. Li T, Elhadi D, Chen G-Q. Co-production of microbial polyhydroxyalkanoates with other chemicals. Metabolic Engineering. 2017 Sep 1;43:29–36.

12. Kumar P, Kim BS. Valorization of polyhydroxyalkanoates production process by co-synthesis of value-added products. Bioresource Technology. 2018 Dec 1;269:544–56.

13. Yadav B, Talan A, Tyagi RD, Drogui P. Concomitant production of value-added products with polyhydroxyalkanoate (PHA) synthesis: A review. Bioresource Technology. 2021 Oct;337:125419.

14. Zhang Y, Li J, Meng J, Sun K, Yan H. A neutral red mediated electro-fermentation system of Clostridium beijerinckii for effective co-production of butanol and hydrogen. Bioresource Technology. 2021 Jul;332:125097.

15. Raj K, Krishnan C. Improved co-production of ethanol and xylitol from low-temperature aqueous ammonia pretreated sugarcane bagasse using two-stage high solids enzymatic hydrolysis and Candida tropicalis. Renewable Energy. 2020 Jun;153:392–403.

16. de Souza Queiroz S, Jofre, dos Santos, Hernández-Pérez, Felipe M Xylitol and ethanol co-production from sugarcane bagasse and straw hemicellulosic hydrolysate supplemented with molasses.

17. Collas F, Kuit W, Clément B, Marchal R, López-Contreras AM, Monot F. Simultaneous production of isopropanol, butanol, ethanol and 2,3-butanediol by Clostridium acetobutylicum ATCC 824 engineered strains. AMB Express. 2012;2(1):45.

18. Julien-Laferrière A, Bulteau L, Parrot D, Marchetti-Spaccamela A, Stougie L, Vinga S, et al. A Combinatorial Algorithm for Microbial Consortia Synthetic Design. Scientific Reports. 2016 Jul 4;6(1):29182.

19. Pharkya P, Burgard AP, Maranas CD. Exploring the Overproduction of Amino Acids Using the Bilevel Optimization Framework OptKnock. Biotechnology and Bioengineering. 2003;84(7):887–99.

20. Kumelj T, Sulheim S, Wentzel A, Almaas E. Predicting Strain Engineering Strategies Using iKS1317: A Genome-Scale Metabolic Model of Streptomyces coelicolor. Biotechnology Journal. 2019;14(4).

21. Choi HS, Lee SY, Kim TY, Woo HM. In Silico Identification of Gene Amplification Targets for Improvement of Lycopene Production. Appl Environ Microbiol. 2010 May 15;76(10):3097–105.

22. Ranganathan S, Suthers PF, Maranas CD. OptForce: An Optimization Procedure for Identifying All Genetic Manipulations Leading to Targeted Overproductions. Price ND, editor. PLoS Comput Biol. 2010 Apr 15;6(4):e1000744.

23. Varma A, Palsson BO. Metabolic Flux Balancing: Basic Concepts, Scientific and Practical Use. Nat Biotechnol. 1994 Oct;12(10):994–8.

24. Kauffman KJ, Prakash P, Edwards JS. Advances in flux balance analysis. Current Opinion in Biotechnology. 2003 Oct 1;14(5):491–6.

25. Orth JD, Thiele I, Palsson BØ. What is flux balance analysis? Nat Biotechnol. 2010 Mar;28(3):245–8.

26. Gudmundsson S, Thiele I. Computationally efficient flux variability analysis. BMC Bioinformatics. 2010 Sep 29;11(1):489.

27. Heirendt L, Arreckx S, Pfau T, Mendoza SN, Richelle A, Heinken A, et al. Creation and analysis of biochemical constraint-based models using the COBRA Toolbox v.3.0. Nat Protoc. 2019 Mar;14(3):639–702.

28. Monk JM, Lloyd CJ, Brunk E, Mih N, Sastry A, King Z, et al. iML1515, a knowledgebase that computes Escherichia coli traits. Nat Biotechnol. 2017 Oct;35(10):904–8.

29. Xu J-Z, Ruan H-Z, Liu L-M, Wang L-P, Zhang W-G. Overexpression of thermostable meso-diaminopimelate dehydrogenase to redirect diaminopimelate pathway for increasing L-lysine production in Escherichia coli. Sci Rep. 2019 Dec;9(1):2423.

30. Liang CY, Xu JL, Xu HJ, Qi W, Zhang Y, Luo W, et al. Gene cloning and characterization of an organic solvent-stimulated β-glucosidase and its application for the co-production of ethanol and succinic acid. Cellulose. 2019 Oct;26(15):8237–48.

31. Zhang X, Jantama K, Shanmugam KT, Ingram LO. Reengineering Escherichia coli for Succinate Production in Mineral Salts Medium. Applied and Environmental Microbiology. 2009 Dec 15;75(24):7807–13.

32. Utrilla J, Gosset G, Martinez A. ATP limitation in a pyruvate formate lyase mutant of Escherichia coli MG1655 increases glycolytic flux to d-lactate. Journal of Industrial Microbiology and Biotechnology. 2009 Aug 1;36(8):1057–62.

33. Bill RM. Playing catch-up with Escherichia coli: using yeast to increase success rates in recombinant protein production experiments.

34. Mo ML, Palsson BØ, Herrgård MJ. Connecting extracellular metabolomic measurements to intracellular flux states in yeast. BMC Syst Biol. 2009;3(1):37.

35. Moxley WC, Eiteman MA. Pyruvate Production by Escherichia coli by Use of Pyruvate Dehydrogenase Variants. Applied and Environmental Microbiology. 87(13):e00487–21.

36. Minois N. Molecular Basis of the ‘Anti-Aging’ Effect of Spermidine and Other Natural Polyamines - A Mini-Review. GER. 2014;60(4):319–26.

37. van Maris AJA, Geertman J-MA, Vermeulen A, Groothuizen MK, Winkler AA, Piper MDW, et al. Directed Evolution of Pyruvate Decarboxylase-Negative Saccharomyces cerevisiae, Yielding a C2-Independent, Glucose-Tolerant, and Pyruvate-Hyperproducing Yeast. Applied and Environmental Microbiology. 2004 Jan 1;70(1):159–66.

38. Zahoor A, Küttner FTF, Blank LM, Ebert BE. Evaluation of pyruvate decarboxylase-negative Saccharomyces cerevisiae strains for the production of succinic acid. Engineering in Life Sciences. 2019;19(10):711–20.

39. Nanda S, Golemi-Kotra D, McDermott JC, Dalai AK, Gökalp I, Kozinski JA. Fermentative production of butanol: Perspectives on synthetic biology. New Biotechnology. 2017 Jul 25;37:210–21.

40. Akhtar J, Idris A, Abd. Aziz R. Recent advances in production of succinic acid from lignocellulosic biomass. Appl Microbiol Biotechnol. 2014 Feb;98(3):987–1000.

41. Xu C, Zhang J, Zhang Y, Guo Y, Xu H, Xu J. Long chain alcohol and succinic acid co-production process based on full utilization of lignocellulosic materials. Current Opinion in Green and Sustainable Chemistry. 2018 Dec 1;14:1–9.

42. Kondo T, Tezuka H, Ishii J, Matsuda F, Ogino C, Kondo A. Genetic engineering to enhance the Ehrlich pathway and alter carbon flux for increased isobutanol production from glucose by Saccharomyces cerevisiae. Journal of Biotechnology. 2012 May 31;159(1):32–7.

43. Dzurendova S, Zimmermann B, Kohler A, Tafintseva V, Slany O, Certik M, et al. Microcultivation and FTIR spectroscopy-based screening revealed a nutrient-induced co-production of high-value metabolites in oleaginous Mucoromycota fungi. PLOS ONE. 2020 Jun 22;15(6):e0234870.

44. Badri A, Raman K, Jayaraman G. Uncovering Novel Pathways for Enhancing Hyaluronan Synthesis in Recombinant Lactococcus lactis: Genome-Scale Metabolic Modeling and Experimental Validation. Processes. 2019 Jun;7(6):343.

45. Srinivasan A, S V, Raman K, Srivastava S. Rational metabolic engineering for enhanced alpha-tocopherol production in Helianthus annuus cell culture. Biochemical Engineering Journal. 2019 Nov;151:107256.

46. Boghigian BA, Armando J, Salas D, Pfeifer BA. Computational identification of gene over-expression targets for metabolic engineering of taxadiene production. Appl Microbiol Biotechnol. 2012 Mar;93(5):2063–73.

47. Kang Z, Gao C, Wang Q, Liu H, Qi Q. A novel strategy for succinate and polyhydroxybutyrate co-production in Escherichia coli. Bioresource Technology. 2010 Oct 1;101(19):7675–8.

